# High-quality *de novo* genome assembly of *Kappaphycus alvarezii* based on both PacBio and HiSeq sequencing

**DOI:** 10.1101/2020.02.15.950402

**Authors:** Shangang Jia, Guoliang Wang, Guiming Liu, Jiangyong Qu, Beilun Zhao, Xinhao Jin, Lei Zhang, Jinlong Yin, Cui Liu, Guangle Shan, Shuangxiu Wu, Lipu Song, Tao Liu, Xumin Wang, Jun Yu

**Affiliations:** College of Life Sciences, Yantai University, Yantai 264005, China; College of Grassland Science and Technology, China Agricultural University, Beijing 100193, China; College of Marine Life Sciences, Ocean University of China, Qingdao 266003, China; CAS Key Laboratory of Genome Sciences and Information, Beijing Key Laboratory of Genome and Precision Medicine Technologies, Beijing Institute of Genomics, Chinese Academy of Sciences, Beijing 100101, China; University of Chinese Academy of Sciences, Beijing 100049, China; Beijing Key Laboratory of Agricultural Genetic Resources and Biotechnology, Beijing Agro-biotechnology Research Center, Beijing Academy of Agriculture and Forestry Sciences, Beijing 100097, China; Key Laboratory of Pratacultural Science, Beijing Municipality 100193, China

**Keywords:** *Kappaphycus alvarezii*, genome assembly, PacBio sequencing, HiSeq sequencing

## Abstract

The red algae *Kappaphycus alvarezii* is the most important aquaculture species in *Kappaphycus*, widely distributed in tropical waters, and it has become the main crop of carrageenan production at present. The mechanisms of adaptation for high temperature, high salinity environments and carbohydrate metabolism may provide an important inspiration for marine algae study. Scientific background knowledge such as genomic data will be also essential to improve disease resistance and production traits of *K. alvarezii*. 43.28 Gb short paired-end reads and 18.52 Gb single-molecule long reads of *K. alvarezii* were generated by Illumina HiSeq platform and Pacbio RSII platform respectively. The *de novo* genome assembly was performed using Falcon_unzip and Canu software, and then improved with Pilon. The final assembled genome (336 Mb) consists of 888 scaffolds with a contig N50 of 849 Kb. Further annotation analyses predicted 21,422 protein-coding genes, with 61.28% functionally annotated. Here we report the draft genome and annotations of *K. alvarezii*, which are valuable resources for future genomic and genetic studies in *Kappaphycus* and other algae.

## Background & Summary

*Kappaphycus alvarezii*, also known as elkhorn sea moss, has the largest individual wet weight in red algae, and is mainly distributed in tropical waters ^1^. They provide important raw materials used for extracting carrageenan, and are large-scale commercially cultivated, mainly in Southeast Asian countries, such as Indonesia, Malaysia, Vietnam and Philippines ^2-4^. Owing to its important economic value as a food source and in the carrageenan industry, *K. alvarezii* cultivation has been introduced into other tropical and subtropical countries ^5^, and the cultivation of the seaweeds *K. alvarezii* and *Eucheuma* spp. has become the most popular in the largest aquaculture production, because κ-Carrageenan as commercial carrageenan applied in food industry is mainly extracted from *K. alvarezii* ^4^. Since in the 1980s *K. alvarezii* was introduced to China, its production is expanded in a large scale ^6,7^.

It is known that red algae with more than 6,000 described species represent the biggest species-rich group in marine macrophytes ^8^. And in evolutionary perspective, red algae are also within the phylogenetic group formed during the endosymbiosis event according to endosymbiosis theory ^9^, and their genes and genomes are crucial for understanding eukaryote evolution. Especially, *K. alvarezii* is ecologically an important component in many marine ecosystems, including rocky intertidal shores and coral reefs. Compared with other unicellular algae and higher land plants, there is a lack of genomic knowledge for *Kappaphycus*. In the macro-algae subclass of Florideophyceae in red algae, the genome of *Chondrus crispus* was firstly published ^10^, whose size is 105 Mb. Therefore, the 336 Mb genome assembly of *K. alvarezii* reported here is effectively promoting the researches in biological metabolism, comparative genomic analysis in algae and eukaryotic evolution, and also potentially provides valuable information for improving economic quality and resistance to environmental changes in aquaculture.

## Methods

### Sample collection and sequencing

*K. alvarezii* strain No.2012020004A provided by Ocean University of China was selected as genomic DNA donor for whole genome sequencing. It was originally from Sulawesi in Indonesia, and cultivated in China by vegetative propagation. To remove the contaminants, the frond (sporophyte) tender tissue was carefully washed in pure water and cut before being immersed in 0.5 g/L I_2_-KI for 15 seconds. And then tissues were washed multiple times and cultivated in sterile sea water at 24°C and 3000 lx for light intensity. The clean frond tissues were used for genomic DNA extraction with the improved CTAB method ^11^, and the library construction was followed.

The pair-end sequencing on Illumina HiSeq platform was performed at Beijing Institute of Genomics, Chinese Academy of Sciences (BIG, CAS) based on the standard protocols. Genomic DNA was fragmented by sonication in Covaris S220 (Woburn, Covaris), and libraries with 300-bp and 500-bp insert size were constructed by using NEBNext® Ultra™ II DNA Library Prep (Ipswich, NEB). The pair-end sequencing was performed, and a total of 214 M reads were generated, i.e. 43.28 Gb raw data, which was about 128-fold coverage of the genome size. At the same time, high-molecular-weight DNA was extracted and 20-kb SMRTbell library was built with size selection protocol on the BluePippin. The *K. alvarezii* genome was sequenced using 16 SMRT cells P6-C4 chemistry on the PacBio RS II platform (at BIG, CAS). The sequencing produced about 18.52 Gb data with an average read length of 10,165 bp, and represented about 55-fold coverage of the genome. All information about sequencing data are shown in Table 1. The raw HiSeq data was filtered using SolexaQA+ software before further analysis ^12^.

**Table 1:**
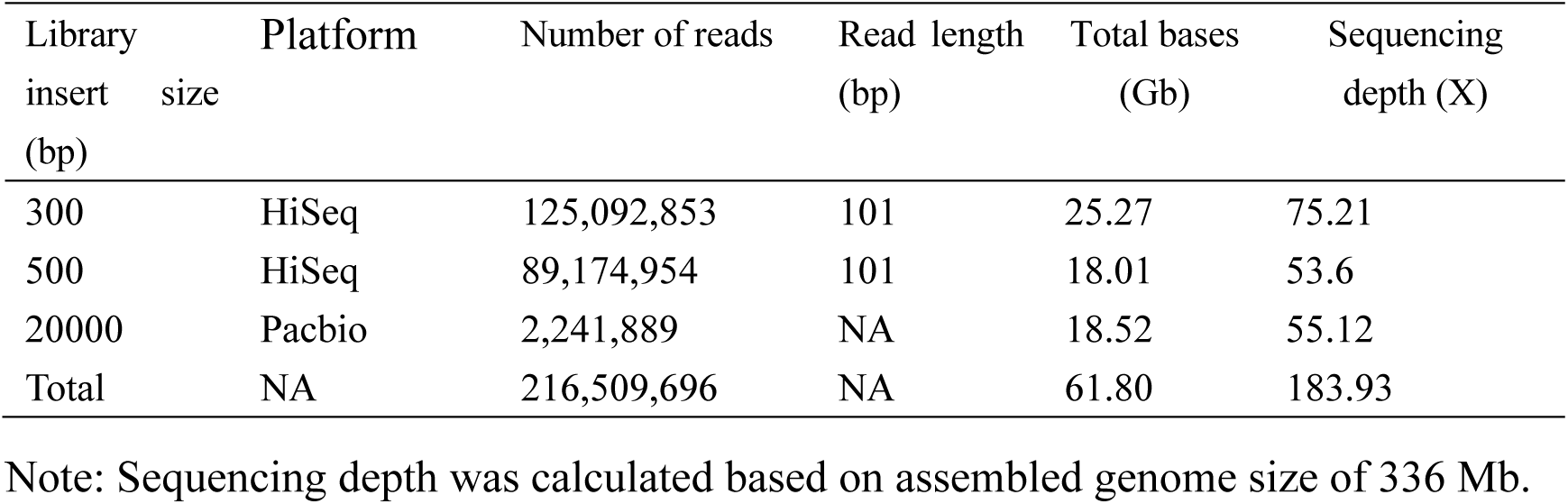
Summary statistics of sequence data in *K. alvarezii* strain No.2012020004A

### *De novo* genome assembly and preliminary evaluation

*De novo* genome assembly of PacBio reads were first performed using Canu with the default parameters to yield the first primary assembly ^13^. And meanwhile, the PacBio reads were assembled into phased diploid assembly using FALCON and FALCON-Unzip, which produced a set of partially phased primary contigs and fully phased haplotigs which represent divergent haplotyes. Then a consensus assembly was generated from the two primary assemblies by canu and FALCON (Fig. 1), by using our locally written Perl scripts. Short reads from Illumina platform were aligned to the assembly using bwa ^14^, followed with duplication removal using Picard tools (http://broadinstitute.github.io/picard/). Pilon was used to do the polish step to correct single insertions and deletions ^15^.

**Figure 1:**
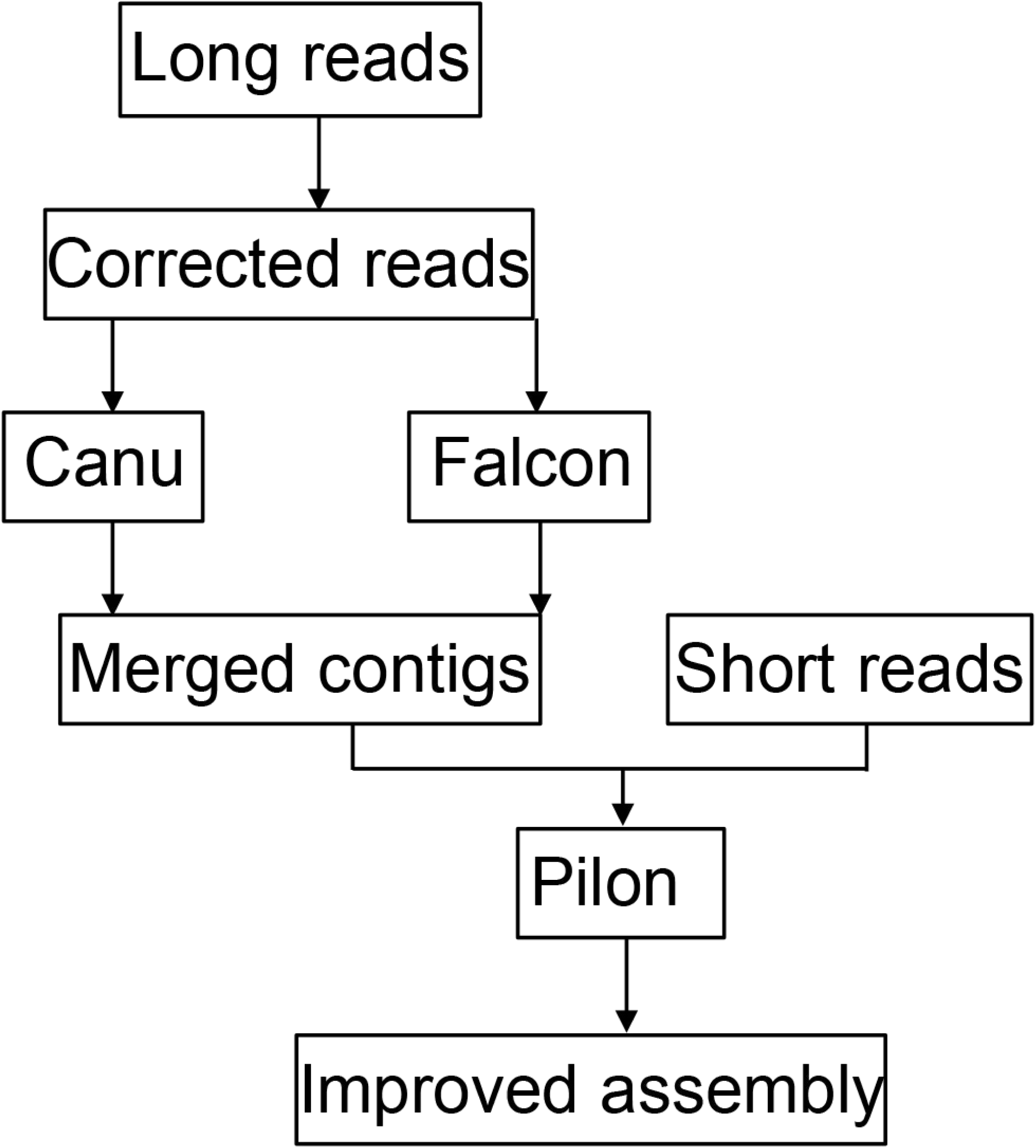
Assembly pipeline for the *K. alvarezii* genome.

We screened all the assembled contigs, and found 11 ones which can be almost 100% mapped to *K. alvarezii* chloroplast complete genome (NCBI accession KU892652.1). Only one contig covered the whole chloroplast genome, and all the 11 chloroplast contigs have been removed out of the assembled contigs. However, we did not find any mitochondrial contigs with a blastn against the complete mitochondrial genome (NCBI accession NC_031814.1). In addition, we tried to filter the bacterial contigs by using blastn against the nt database, and none were found with identity > 90%. Finally, this led to a genome assembly of 336,052,185 Mb with a contig N50 size of 849,038 bp, and the quality of this assembly is high enough for the downstream analysis (Table 2).

**Table 2:**
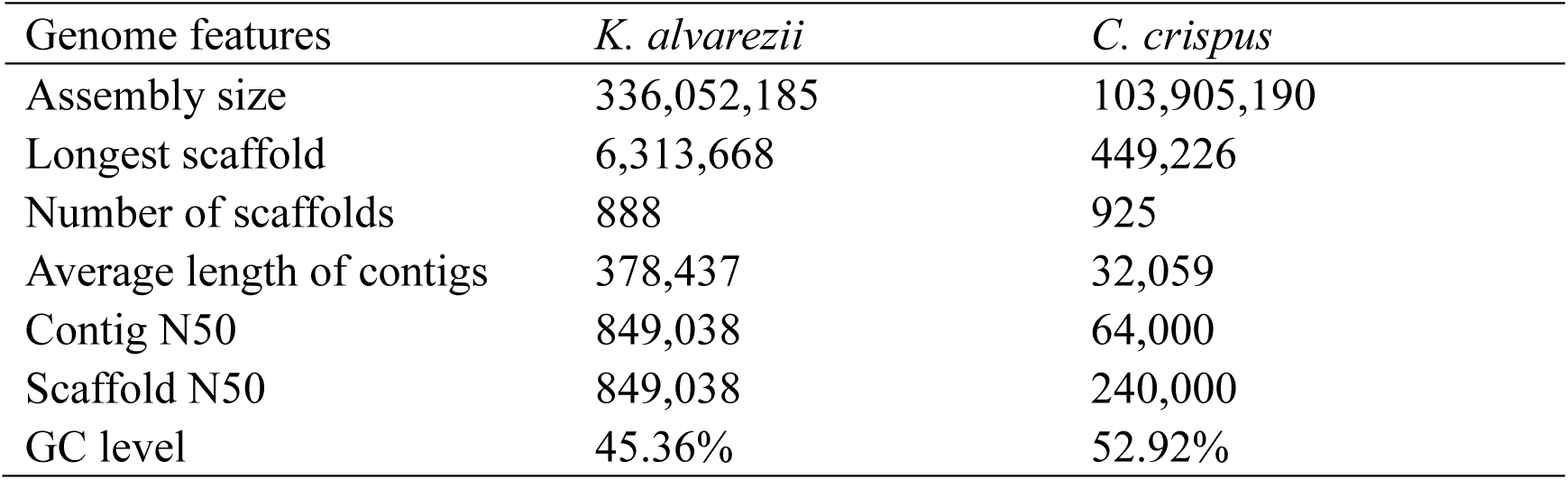
Summary statistics of the genome assemblies in *K. alvarezii and C. crispus*

| Genome features | *K. alvarezii* | *C. crispus* |
| --- | --- | --- |
| Assembly size | 336,052,185 | 103,905,190 |
| Longest scaffold | 6,313,668 | 449,226 |
| Number of scaffolds | 888 | 925 |
| Average length of contigs | 378,437 | 32,059 |
| Contig N50 | 849,038 | 64,000 |
| Scaffold N50 | 849,038 | 240,000 |
| GC level | 45.36% | 52.92% |

Furthermore, to evaluate the completeness of the assembly, a set of ultra-conserved core eukaryotic genes identified by CEGMA were mapped to the assembled genome using CEGMA ^16^ and BUSCO ^17^, which quantitatively assess genome completeness using evolutionarily informed expectations of gene content. CEGMA assessment showed that our assembly captured 228 (91.94%) of the 248 ultra-conserved core eukaryotic genes, of which 214 (86.29%) were complete (Table S1). BUSCO assessment showed that the assembly captured 264 (87.13%) of the 303 ultra-conserved core eukaryotic genes (eukaryota_odb9), of which 259 (85.5%) were complete, while 10.2% were considered missing in the assembly (Table 3). It was comparable with the results of *C. crispus* assembly.

**Table 3:** Summarized benchmarking in BUSCO notation for the assembly

|  | *K. alvarezii* |  | *C. crispus* |  |
| --- | --- | --- | --- | --- |
|  | Number | Percent | Number | Percent |
| Complete BUSCOs (C) | 259 | 85.50% | 263 | 86.80% |
| Complete and single-copy BUSCOs (S) | 176 | 58.10% | 254 | 83.80% |
| Complete and duplicated BUSCOs (D) | 83 | 27.40% | 9 | 3.00% |
| Fragmented BUSCOs (F) | 13 | 4.30% | 10 | 3.30% |
| Missing BUSCOs (M) | 31 | 10.20% | 30 | 9.90% |

Note: totally 303 BUSCO groups were searched. BUSCO was run in mode genome, the lineage dataset is eukaryota\_odb9.

### Repeat annotation in the genome assembly

We used two methods to identify the repeat contents in *K. alvarezii* genome, i.e. homology-based one and *de novo* prediction. The homology-based analysis was performed by RepeatMasker (http://www.repeatmasker.org/) using the repetitive database of RepBase ^18^. In *de novo* prediction, RepeatMasker (version 3.3.0) was used to identify transposable repeats in the genome with a *de novo* repeat library constructed by RepeatModeler v1.0.8 (http://www.repeatmasker.org/RepeatModeler/). Blast searches were followed to classify those elements, at the DNA level: E-value <=1e-5, identity percent >=50%, alignment coverage>=50%, and the minimal matching length >=80bp; and at the protein level: E-value <=1e-4, identity percent >=30%, alignment coverage>=30%, and the minimal matching length >=30 amino acids. In conclusion, more than 179 million bases were found as interspersed repeats in the *K. alvarezii* genome, covered about 53.35% of the genome size (Table 4). The most abundant transposable elements were LTR elements (27.58%), LINES (8.61%), and DNA transposons (5.75%).

**Table 4:** Summary statistics of annotated repeats in the assembly

|  | Number of elements | Length occupied (bp) | Percentage of sequence |
| --- | --- | --- | --- |
| SINEs | 355 | 76,627 | 0.02% |
| LINEs | 65,095 | 28,994,744 | 8.61% |
| LTR elements | 67,843 | 92,875,333 | 27.58% |
| DNA elements | 36,440 | 19,350,040 | 5.75% |
| Unclassified | 121,683 | 38,353,941 | 11.39% |
| Total interspersed repeats | 291,416 | 179,650,685 | 53.35% |

Note: most repeats fragmented by insertions or deletions have been counted as one element.

### Gene prediction and functional annotation

Three approaches for gene model prediction, i.e. homology detection, expression-evidence-based predictions and ab initio gene predictions, were combined to get consensus gene structures. To identify homology patterns in *K. alvarezii*, the BLASTX ^19^ search was conducted against the NCBI non-redundant protein database with E-value <10^−5^, and then the proteins were aligned for their gene structure by GeneWise ^20^, and introns and frameshifting errors were identified. For expression evidences, published ESTs, transcripts and RNA-seq datasets were aligned to the genome. AUGUSTUS was used for *ab initio* gene prediction ^21^ after that repeated elements in the nuclear genome were masked by RepeatMasker. Gene model parameters for the programs were trained based on long transcripts and known *Kappaphycus* genes. And then, all these *de novo* gene predictions, homolog-based methods and RNA-seq data were combined to determine the consensus gene sequences using EVidenceModeler (EVM) ^22^, and PASA was used to update the EVM consensus predictions by adding UTR annotations, merging genes, splitting genes, boundary adjustments ^23^. It resulted in 21,422 protein-coding gene models. The gene length distribution, coding sequences (CDS), exons, introns, and the distribution of exon number per gene were shown in Table 5. Totally 254 contigs do not contain protein-coding genes, i.e. 12,285,700 bp in length and 3.6% of the whole assembly.

**Table 5:** Summary statistics of gene structure

|  | *K. alvarezii* | *C. crispus* |
| --- | --- | --- |
| Protein-coding loci | 21,422 | 9,606 |
| Average length of transcript | 1089.12 | - |
| Average length of cds | 981 | 1,080 |
| Average length of exon | 500.90 | 789 |
| Average number of exon | 1.98 | 1.32 |
| Average length of intron | 422.03 | 123 |

For functional assignment and annotation, the BLAST search of gene models was carried out against NR, Swissprot and TrEMBL protein database ^24^ with E-value <10^−5^. While InterProScan program ^25^ was used to perform functional classification of Gene Ontology (GO) of the genes, and also generate family information from Interpro. Pathway analysis was performed using the Kyoto Encyclopedia of Genes and Genomes (KEGG) annotation service KAAS with the default bitscore threshold of 60 ^26^. Totally 13,011 proteins were annotated, i.e. 60.7% of all predicted proteins (Table 6 & Table S2). The all-vs-all BLAST search against genes themselves identified the distribution of gene copies in the whole genome based on the identity (Fig. S1), and it showed that the most genes were with one or two copies for 100% identity, and more homologs were identified with smaller identity.

**Table 6:** Statistics for functional annotation

|  | Number | Percent (%) |
| --- | --- | --- |
| Nr | 12666 | 59.69% |
| Swissprot | 8145 | 38.78% |
| TrEMBL | 12705 | 59.86% |
| InterPro | 9202 | 43.65% |
| KEGG | 4,448 | 19.72% |
| GO | 9,642 | 42.74% |
| Total | 13,011 | 61.28% |

Furthermore, we selected 22 conserved genes and downloaded their homologous sequences from 14 plant species, including spermatophyte, Bryophyta, Charophyta, Chlorophyta, Glaucophyta, and Rhodophyta. We built a phylogenetic tree based on these homologous sequences, and found that *K. alvarezii* was placed with a close position to *C. crispus* (Fig. 2), which is consistent with the result in the Nr database search (Fig. 3).

**Figure 2:**
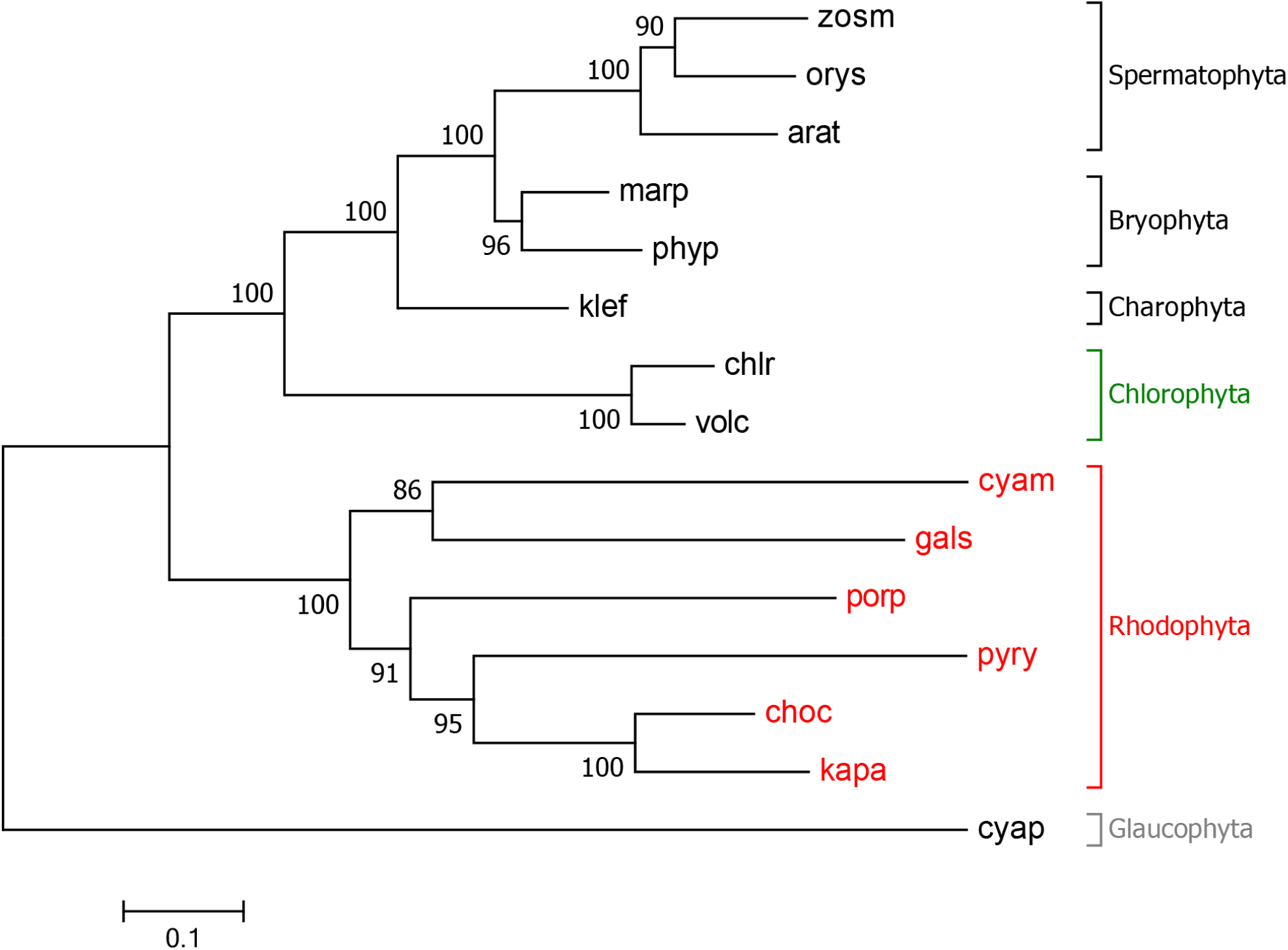
Molecular Phylogenetic analysis by Maximum Likelihood method, inferred by using the Maximum Likelihood method based on the Le_Gascuel_2008 model (LG+G). *Arabidopsis thaliana*, arat; *Chlamydomonas reinhardtii*, chlr; *Chondrus crispus*, choc; *Cyanidioschyzon merolae*, cyam; *Cyanophora paradoxa*, cyap; *Galdieria sulphuraria*, gals; *Kappaphycus alvarezii*, kapa; *Klebsormidium flaccidum*, klef; *Marchantia polymorpha*, marp; *Oryza sativa*, orys; *Physcomitrella patens*, phyp; *Porphyridium purpureum*, porp; *Pyropia yezoensis*, pyry; *Volvox carteri*, volc; *Zostera marina*, zosm.

**Figure 3:**
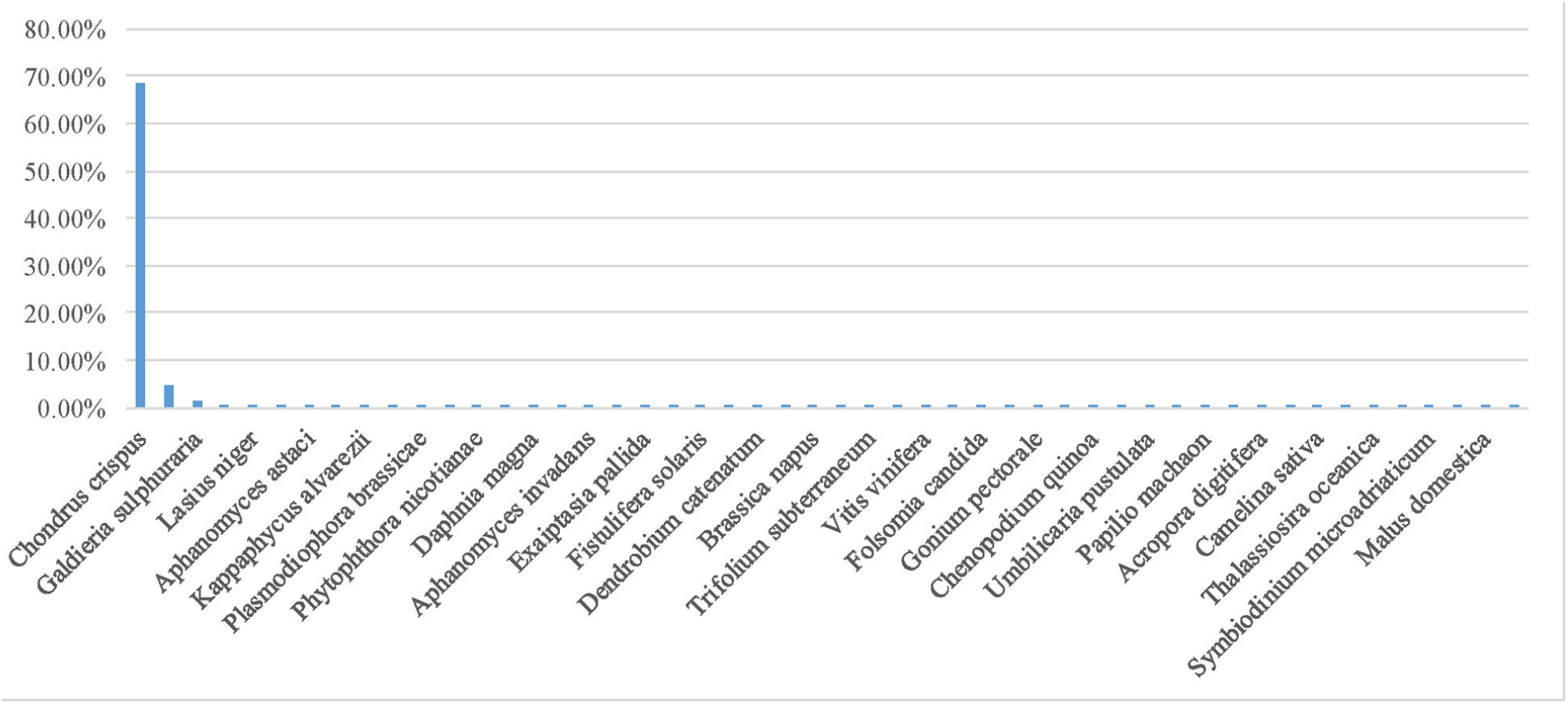
Blast annotation against the NCBI nr database.

## Data Records

All of the raw reads have been deposited at SRA under the accession numbers of SRP101845 and SRP128943. This whole genome shotgun project has been deposited at DDBJ/ENA/GenBank under the accession NADL00000000. The version described in this paper is NADL01000000.2. The raw sequence data has also been deposited in the Genome Sequence Archive ^27^ in BIG Data Center ^28^, Beijing Institute of Genomics (BIG), Chinese Academy of Sciences, under accession numbers PRJCA000373 that are publicly accessible at http://bigd.big.ac.cn/gsa.

## Technical Validation

Genome size was estimated by the k-mer method using Jellyfish and gce program ^29^. K-mer analysis was performed by using 34.15 Gb clean sequences from 300 and 500 bp insert size libraries, and the estimated genome size of *K. alvarezii* was 334,905,000 bp. Furthermore, it is shown in a previous study that there are ten chromosomes (n = 10) in *K. alvarezii* nucleus, and the g/2C genome size based on the cytophotometry was estimated to be 0.28∼0.32 pg ^30^. The genome of *K. alvarezii* was therefore extrapolated to be 273.8∼313 Mb (0.978 x 10^9^ bp/pg) ^31^, which is consistent with the genome assembly in this study.

Furthermore, the assembled contigs were evaluated based on the following analysis. Firstly, the coverage peaks for 17 kmer were about 65X and 35X for HiSeq and PacBio reads respectively (Fig. S2A and B), and only one peak was found for 17-, 25- and 30-kmer (Fig. S2C), which suggested a reliable assembly. Secondly, we did BLAST alignment of the assembled contigs against NCBI Nr database, and found the majority was with the hits to *C. crispus*, a species of red algae (Fig. 3). Finally, the depths of HiSeq and Pacbio reads were shown a relatively stable distribution across the assembled contigs, and it suggested no severe bias for both the sequencing methods (Fig. S3).

It was reported that the three second components (fast, intermediate, and slow) in the DNA reassociation kinetic analysis corresponded to the highly repetitive sequences (12%), mid-repetitive sequences (38%) and unique sequences (50%) ^30^, and our repeat ratio of 53.35% further confirmed that almost half of the *K. alvarezii* genome is not unique.

## Usage Notes

We report the first genome sequencing, assembly, and annotation of the red alga *K. alvarezii*. The assembled draft genome will provide a valuable genomic resource for the study of essential genes, especially Carrageenan and other useful polysaccharides; for the alignment of sequencing reads, for example, RNA-seq and low-coverage genome resequencing. And the well-annotated gene sequences are also helpful to conduct more comprehensive evolution analysis of genes in Florideophyceae algae, and understand the genomic evolution in algae.

## Code Availability

Software used for read preprocessing, genome assembly and annotation is described in the Methods section together with the versions used.

## Acknowledgements

We thank the faculty and staff in the BIG of CAS, who contributed to the sequencing of the genome, and the Culture Collection of Seaweed at the Ocean University of China, for providing *K. alvarezii*. This project was supported by the China-ASEAN Maritime Cooperation Fund and Top Talent Program of The Yantai University.

## Authors’ contributions

TL, XW and JY conceived the project. XJ, LZ and CL provided the samples. GW, SJ and GL performed genome assembly, repeat annotation, gene prediction, gene function annotation and other analysis. BZ, JY, GS, LS, and SW were involved in the experiments and analysis. SJ and GW wrote and revised the manuscript. All authors read and approved the final manuscript.

## Supplemental materials

**Figure S1:** The frequency of self-blast alignments of genes, multiple hits for each query were shown in different colors, sorted by blast scores.

**Figure S2:** K-mer distribution in the *K. alvarezii* genome. In A for HiSeq and B for PacBio, the x-axis is frequency (depth) of 17 k-mer; the y-axis is the proportion which represents the frequency at that depth divide by the total frequency of all the depth. C, comparison of 17, 25, and 30 k-mer.

**Figure S3:** Sequencing depth of the contigs, calculated respectively from HiSeq (A) and PacBio (B) data, with 20 kb window size. Contigs larger than 1 Mb were selected for calculation.

**Table S1:** Statistics of the completeness of the genome based on CEGMA.

**Table S2:** Annotation of all genes in the assembly.

