## Supplementary figures and images for "High-quality *de novo* genome assembly of *Kappaphycus alvarezii* based on both PacBio and HiSeq sequencing"

### Figure S1

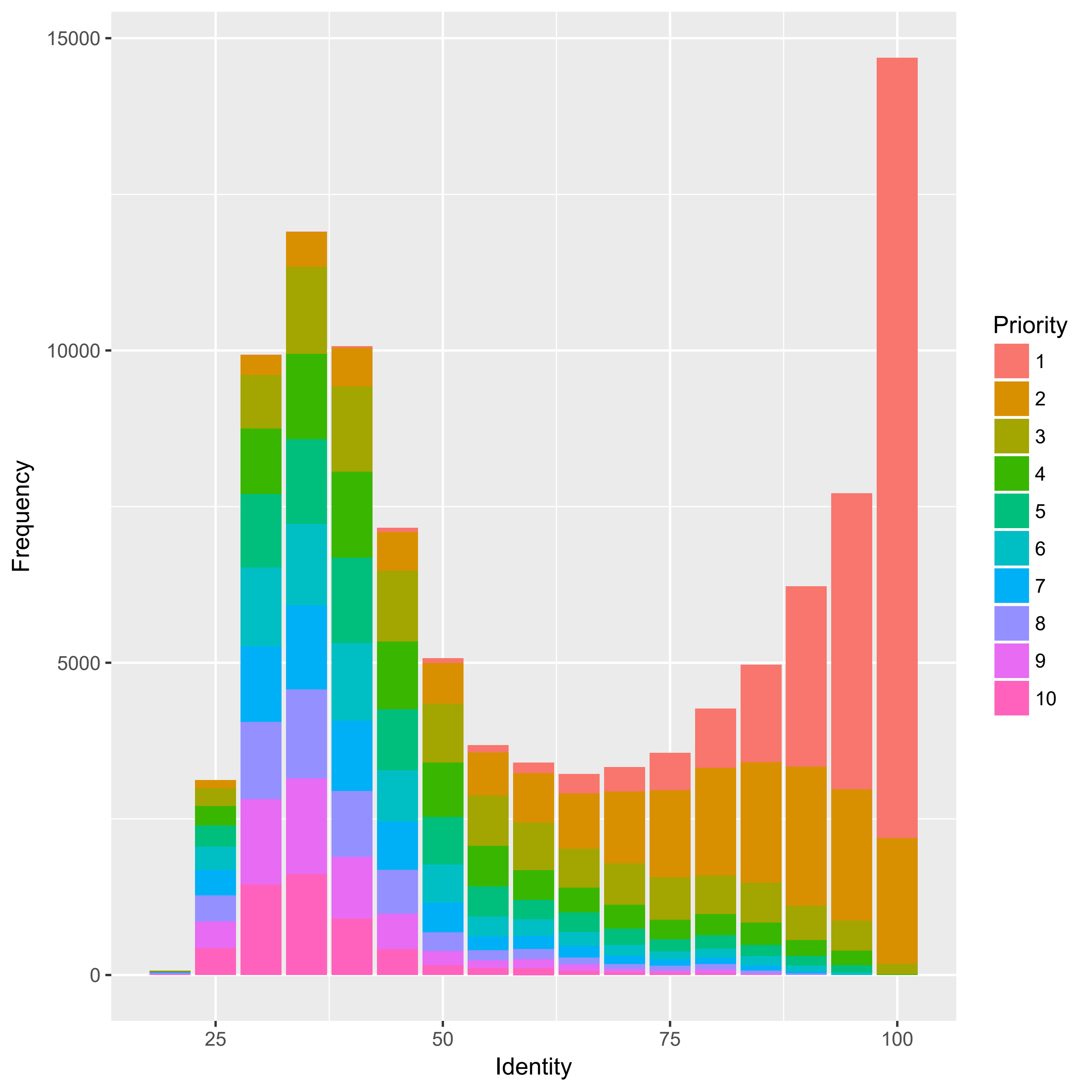

### Figure S2

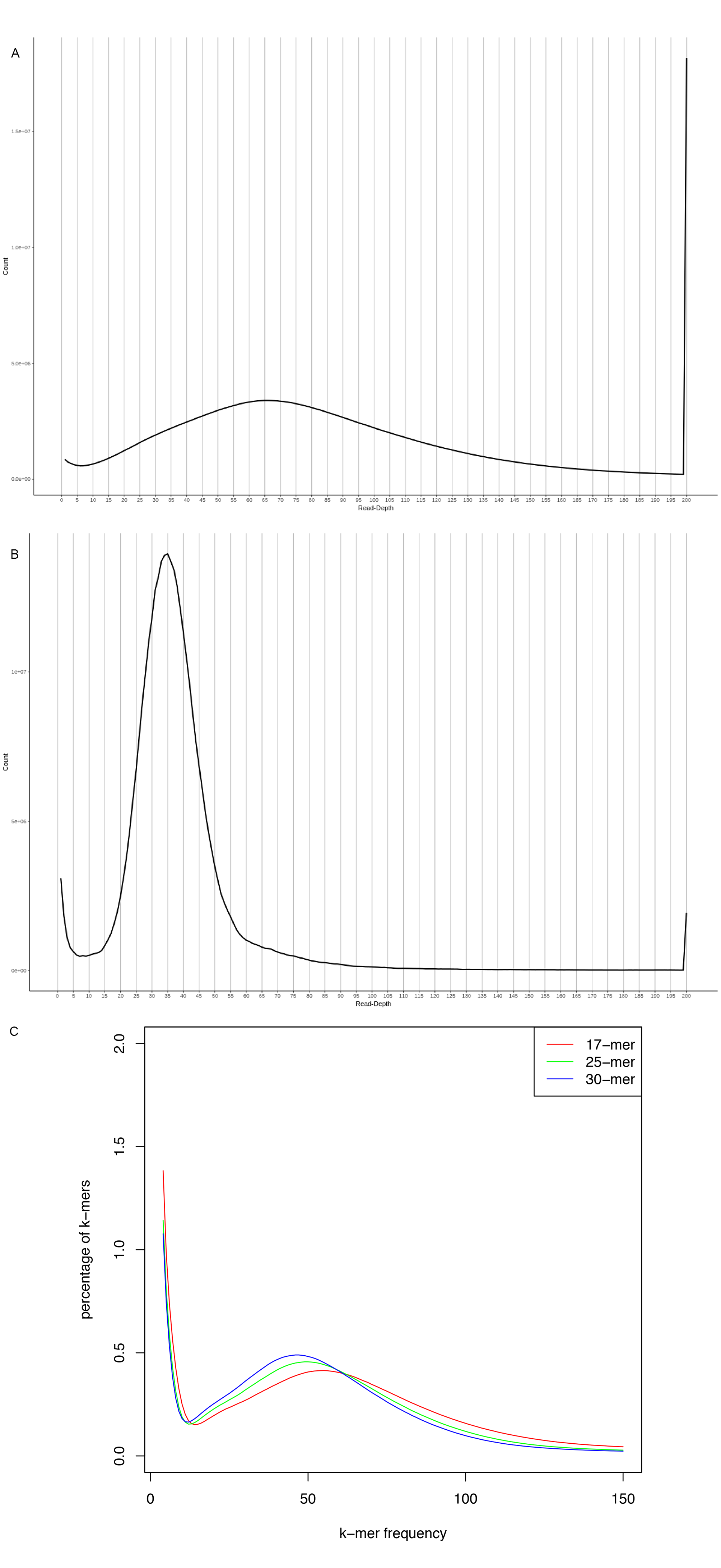

### Figure S3

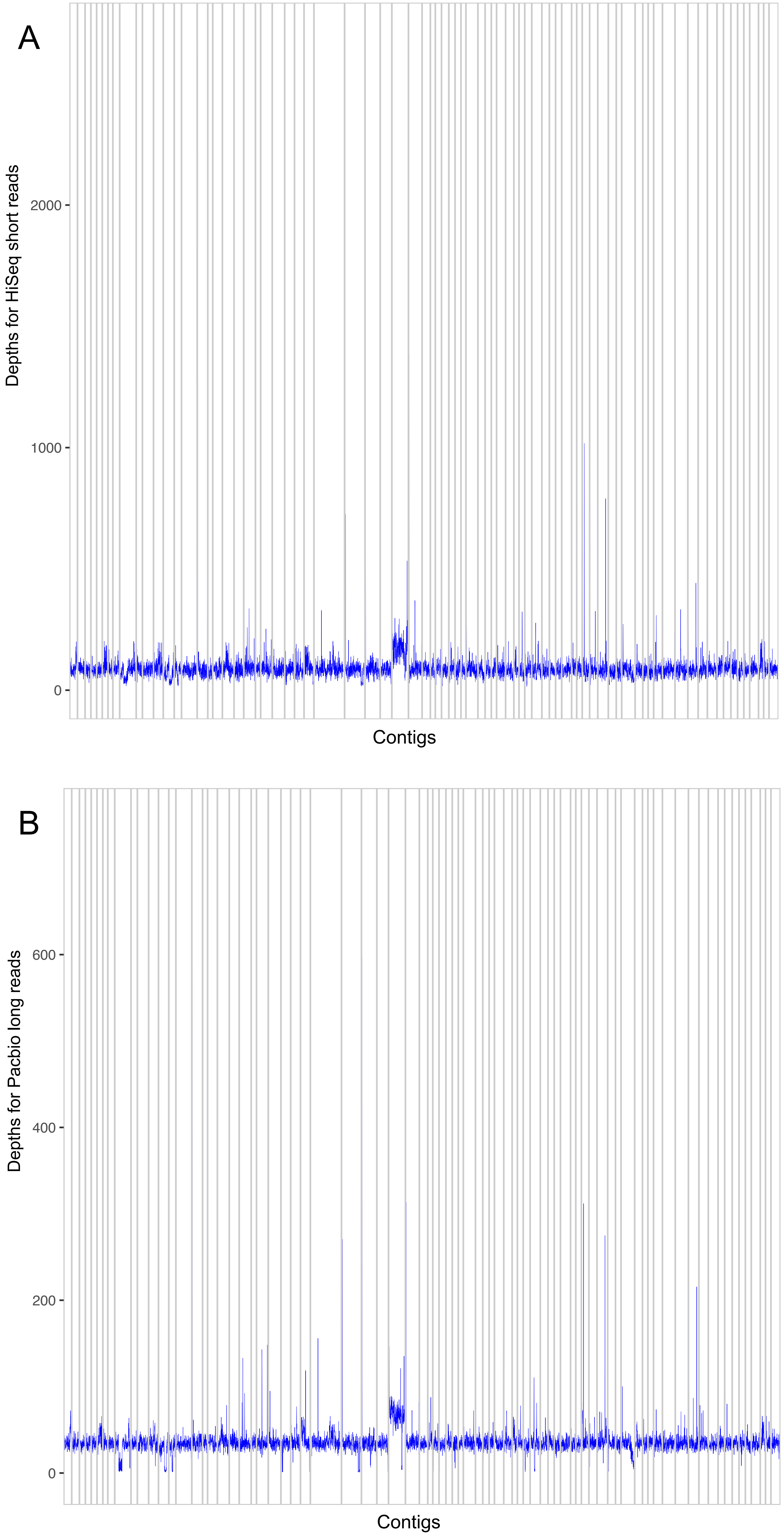
